# Comparing aperiodic brain activity between eyes open rest and dynamic visual input using magnetoencephalography

**DOI:** 10.64898/2026.03.28.714956

**Authors:** Tzu-Yu Hsu, Ko-Ping Chou, Yi-Ju Liu, Niall W. Duncan

## Abstract

Inscapes is a low demand abstract animation used as an alternative to eyes open rest in neuroimaging studies, particularly with pediatric and clinical populations prone to head motion. Although prior work has established that functional connectivity patterns during Inscapes closely resemble those during rest, no study has examined whether the two conditions differ in aperiodic neural activity, a broadband feature of the power spectrum linked to excitation/inhibition balance. Here we used magnetoencephalography (MEG) in 54 healthy adults to compare spectrally parameterised aperiodic and periodic measures between eyes open rest and Inscapes viewing (visual component only, without audio). At the sensor level, both the aperiodic exponent and offset were significantly higher during rest than during Inscapes across widespread frontoparietal and occipital distributions in both magnetometers and gradiometers. Source level analyses at both the parcellation and vertex levels largely supported these patterns. The pericalcarine cortex was a notable exception, where both aperiodic measures were higher during Inscapes than during rest, indicating a regionally specific reversal in primary visual cortex. These results demonstrate that Inscapes and eyes open rest produce distinct aperiodic spectral profiles, indicating that the two conditions are not interchangeable for analyses involving broadband spectral dynamics or excitation/inhibition balance estimation.

## Introduction

Inscapes is a computer-generated animation featuring slowly evolving abstract shapes accompanied by a pentatonic piano score. This was originally developed to help address the influence of head motion and participant drowsiness on resting state neuroimaging data quality (Duncan & Northoff, 2012), particularly in young children and clinical populations (Vanderwal et al., 2015). The stimulus is nonsocial, nonverbal, and contains no narrative arc or scene cuts, thereby minimizing cognitive load while providing sufficient visual engagement to maintain wakefulness and reduce motion. In the original fMRI validation, both Inscapes and a narrative movie clip reduced head motion and self-reported sleep relative to eyes open rest, while functional connectivity patterns during Inscapes more closely resembled rest than those during the narrative movie (Vanderwal et al., 2015). A subsequent study further demonstrated that movie viewing, including Inscapes, enhanced the detection of individually unique functional connectivity signatures relative to rest (Vanderwal et al., 2017). In principle, these properties therefore position Inscapes as a low demand intermediate condition that improves data quality while preserving brain activity properties of interest.

The paradigm has since been adopted in studies involving various participant populations across a range of neuroimaging techniques. For example, Inscapes has been used with the aim of reducing head motion in fMRI research involving infants (Copeland et al., 2022; Frew et al., 2022; Vartiainen et al., 2025) and children with autism spectrum disorder (ASD) (Gabrielsen et al., 2018), as well as in multiple studies with adult participants (e.g. (Kröll et al., 2023; Telesford et al., 2023)).

Similarly, an MEG validation study has confirmed that head motion was reduced when using Inscapes with typically developing children and those with ASD or attention deficit/hyperactivity disorder (ADHD), with comparable paradigm-induced modulations in oscillatory power and functional connectivity observed across diagnostic groups (Vandewouw et al., 2021). The paradigm has also been incorporated into neurostimulation protocols for major depressive disorder and ASD to stabilise connectivity measures across sessions (Plewnia et al., 2021; Prillinger et al., 2021), and into substance use research to minimise motion confounds during neurofeedback and pharmacological challenge designs (Chung et al., 2023; Kirsch et al., 2023). Further validation in functional near-infrared spectroscopy (fNIRS) demonstrated that prefrontal connectivity during Inscapes closely resembled that during a fixation condition and predicted executive function performance in preschoolers and young adults (Eng et al., 2025). Collectively, these studies establish Inscapes as a robust tool to assist in the acquisition of high-quality neuroimaging data across ages, clinical conditions, and imaging modalities.

Despite this growing adoption, existing comparisons between Inscapes and eyes open rest have primarily focused on functional connectivity metrics or total spectral power within canonical frequency bands (Vanderwal et al., 2015, 2017; Vandewouw et al., 2021). However, electrophysiological power spectra comprise not only periodic oscillations but also an aperiodic component following a 1/f distribution (Donoghue et al., 2020; He, 2014). The slope of this 1/f distribution, often referred to as the aperiodic exponent, has been proposed as a proxy for the balance between neural excitation and inhibition (Gao et al., 2017), and is sensitive to cognitive state (Waschke et al., 2021), developmental maturation (Green et al., 2025; Hill et al., 2022; Voytek et al., 2015), and various clinical conditions (e.g., (Akbarian et al., 2025; Kopf et al., 2024; Monchy et al., 2024)). Although recent work has used aperiodic parameters extracted during Inscapes viewing to examine developmental and group differences (Green et al., 2025; Reichelt et al., 2026; Vandewouw et al., 2024), whether aperiodic spectral parameters differ between Inscapes and eyes open rest has not been directly compared in healthy adults. This leaves open questions as to the effect of the video stimulus itself upon the aperiodic component of MEG signals. Such effects would be important considerations for the interpretation of results based upon data acquired using Inscapes.

The present study addresses this gap by using MEG and spectral parameterisation (Donoghue et al., 2020) to compare aperiodic neural activity between eyes open rest and Inscapes viewing. We extracted the aperiodic exponent and offset as primary outcome measures across the whole brain, with alpha band power extracted as a secondary measure. If Inscapes induces an equivalent neural state to eyes open rest, no significant differences in aperiodic parameters would be expected.

Conversely, if the sustained visual stimulation from the Inscapes video modulates the excitation-to-inhibition balance or broadband neural activity, condition-specific changes in the aperiodic exponent or offset should emerge. The nature of any such change could inform interpretations in future research.

## Methods

### Participants

Fifty-six healthy adults were recruited for the MEG experiment (29 females; mean age = 26.6 ± 4.6 years). All participants had normal or corrected-to-normal vision and no self-reported psychiatric diagnoses, neurological disorders, or colour blindness.

Two participants were excluded from further analysis due to insufficient trials remaining after artifact rejection during preprocessing, yielding a final sample of 54 participants.

### Ethics and Compensation

The experimental protocol was approved by the Taipei Medical University Institutional Review Board (IRB N202003129). Written informed consent was obtained from all participants prior to participation, and all participants received financial compensation (1500 NTD, approximately 46 USD) for their time.

### Procedure

All recordings took place in a magnetically and electrically shielded room. Participants sat upright with their head resting against the rear of the MEG dewar. Visual stimuli were back-projected onto a screen positioned 120 cm in front of the participant using a Panasonic projector (1,024 × 768 pixels; 60 Hz refresh rate).

Before entering the scanner, participants completed a set of questionnaires related to other aims of the broader project. They were then given verbal instructions for each condition and reminded to remain as still as possible throughout the recording session. Two seven-minute conditions were administered: eyes open rest (EO) and Inscapes. In the EO condition, a central fixation cross was displayed and participants were asked to let their thoughts wander freely without dwelling on any particular topic. In the Inscapes condition, participants passively viewed the animation and were asked to stay relaxed and awake. Only the visual component of Inscapes was presented; the accompanying audio track was not included.

### MEG/MRI Acquisition

Neuromagnetic signals were acquired with a whole-head 306-channel Triux system (Elekta Neuromag), housed at the Imaging Center for Integrated Body, Mind, and Culture Research, National Taiwan University, Taipei, Taiwan. The sensor array consisted of 102 magnetometers and 204 orthogonal planar gradiometers.

Continuous data were sampled at 1,000 Hz with an online bandpass filter of 0.03–330 Hz. Prior to recording, the head coordinate system was established by digitising three anatomical fiducials (nasion, left and right preauricular points) with a Polhemus Fastrak electromagnetic digitiser (Colchester, VT, United States).

Continuous head position monitoring was enabled by four head position indicator (HPI) coils attached to the left and right forehead and mastoid areas. To facilitate subsequent coregistration with structural MRI data, more than 100 additional points on the scalp and face were digitised for each participant. Ocular artifacts were monitored via vertical electrooculogram (VEOG) electrodes positioned above and below the right eye and horizontal electrooculogram (HEOG) electrodes placed at the outer canthi of both eyes. Cardiac activity was recorded using MEG-compatible electrodes affixed to the left and right wrists.

Structural brain images were obtained in a separate MRI session following MEG data collection. A high-resolution T1-weighted anatomical scan was acquired for each participant on a 3T Siemens MAGNETOM Prisma scanner equipped with a 64-channel head coil, located at the same imaging center. Images were collected using a magnetisation-prepared rapid acquisition gradient echo (MPRAGE) sequence with an isotropic voxel size of 0.9 mm, with 208 slices and a field of view of 256 × 256 mm.

### MRI Preprocessing

Each participant’s T1-weighted structural image was processed using the FreeSurfer *recon-all* pipeline (version 7.4.1)(Fischl, 2012), which performs intensity normalization, skull stripping, white matter segmentation, cortical surface reconstruction, and parcellation. Boundary element model (BEM) surfaces were then generated from the FreeSurfer reconstruction using the watershed algorithm implemented in MNE Python (Gramfort et al., 2013, 2014). Scalp surfaces were additionally created for coregistration purposes. Coregistration between the MEG digitisation points and the reconstructed head surfaces was performed interactively using the MNE coregistration graphical interface, aligning the three fiducial landmarks and iteratively fitting the digitised scalp points to the scalp surface.

### MEG Preprocessing

Raw MEG data were organized into BIDS format (Appelhoff et al., 2019) and processed using MNE Python (Gramfort et al., 2013). Spatiotemporal signal space separation (tSSS) was applied via Maxwell filtering to suppress external and nearby interference, with automatic bad channel detection and reconstruction using system-specific fine calibration and crosstalk correction files (Taulu & Simola, 2006). Head positions were aligned across conditions to the first recording run within each participant. The tSSS-processed data were downsampled to 200 Hz and bandpass-filtered between 1 and 30 Hz for independent component analysis (ICA). A 30-component FastICA decomposition was computed, and components reflecting cardiac and ocular artifacts were identified by visual inspection and removed. The ICA solution was then applied to the original tSSS-processed data to preserve full spectral content.

### Spectral Parameterization

For sensor-level analysis, ICA-cleaned data from each condition were cropped to the final 300 seconds, filtered between 0.1 and 55 Hz, and further cleaned with signal space projections for cardiac and ocular artifacts. The continuous data were segmented into four-second epochs, with amplitude-based rejection applied (gradiometers: 8 × 10⁻¹⁰ T/m; magnetometers: 8 × 10⁻¹² T); a minimum of 10 clean epochs per condition was required. Power spectral density (PSD) was estimated per epoch using Welch’s method with 50% overlap and averaged across epochs using the median. The SpecParam algorithm (Donoghue et al., 2020) was applied to each magnetometer and gradiometer’s averaged PSD to decompose the spectrum into aperiodic and periodic components over a 5 to 50 Hz range (fixed aperiodic mode; peak width limits: 1–6 Hz; maximum 6 peaks; minimum peak height: 0.15; R² threshold > 0.85). The aperiodic exponent and offset were extracted as primary outcome variables, with alpha band (8–13 Hz) center frequency and power extracted as secondary measures. Only participants with valid fits in both conditions were retained.

### Source Level Analysis

Cortical source spaces were constructed on individual FreeSurfer reconstructions using an octahedron-6 subdivision (approximately 4,098 vertices per hemisphere), with single-shell boundary element models (BEM; conductivity = 0.3 S/m). Forward solutions were computed once per participant and reused across conditions. A shared inverse operator was derived from noise covariance pooled across both conditions (loose = 0.2; depth = 0.8), and source time courses were estimated using dynamic statistical parametric mapping (dSPM; λ² = 1/9, normal orientation). For parcellation-level analysis,label time courses were extracted using the Desikan Killiany atlas with PCA flip sign-consistent signal extraction. For vertex-level analysis, PSD was estimated at each vertex using multitaper spectral estimation (bandwidth = 2.0 Hz) and SpecParam fitting was then applied. Results were morphed to the fsaverage6 template using the same SpecParam parameters as at the sensor level.

### Statistical Analysis

Condition differences between eyes open rest and Inscapes were assessed using cluster-based permutation paired t-tests (Maris & Oostenveld, 2007). At the sensor level, the paired difference (rest minus Inscapes) for each spectral parameter was tested across all gradiometers and magnetometers separately. Clusters of spatially contiguous sensors exceeding a two-tailed threshold (*p* < 0.05) were identified using the sensor adjacency structure, and cluster-level significance was evaluated with 5000 sign-flip permutations. At the source level, vertex-wise cluster-based permutation tests were conducted on the fsaverage surface using spatial adjacency, with 5000 permutations and family-wise error rate controlled at *α* = 0.05. For the parcellation-level analysis, condition differences were assessed using paired t-tests across 68 Desikan-Killiany atlas labels, with false discovery rate (FDR) correction applied across labels separately for each spectral parameter. Participants with more than 50% missing labels due to poor spectral fits were excluded from each respective analysis.

## Results

### Aperiodic Exponent

Aperiodic and periodic spectral components were estimated separately for magnetometers and gradiometers. For magnetometers (Figure 1, first row), cluster-based permutation tests revealed a significant positive cluster for the aperiodic exponent spanning 88 sensors (cluster statistic = 367.40, *p* < .001), indicating steeper aperiodic exponents during eyes open rest than during Inscapes across a broad frontoparietal and occipital distribution. Gradiometers (Figure 1, second row) similarly showed a large significant positive cluster spanning 138 sensors (cluster statistic = 549.56, *p* < .001), with higher exponent values during eyes open rest across a widespread scalp distribution.

**Figure 1.**
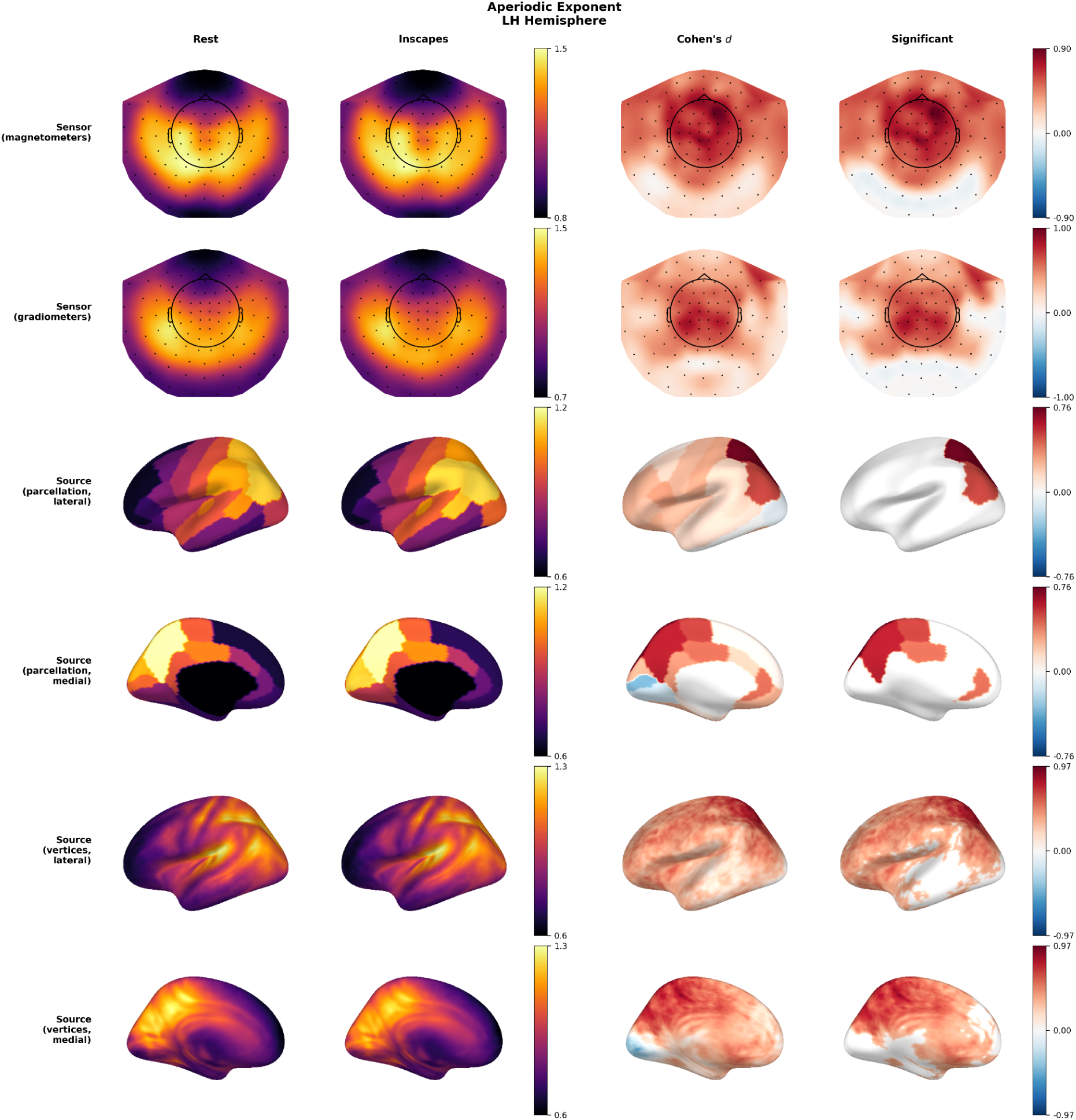
Multi level comparison of aperiodic exponents between eyes open rest and Inscapes, left hemisphere. Rows display results for magnetometers (first row), gradiometers (second), source-level parcellation on the Desikan–Killiany atlas in lateral view (third) and medial view (fourth), and source-level vertices on the fsaverage6 surface in lateral view (fifth) and medial view (sixth). Columns show the group-mean value for the rest condition (first), the Inscapes condition (second), paired Cohen’s d effect size for the rest minus Inscapes contrast (third), and the statistically significant regions with non-significant areas masked to zero (fourth). The number of significant sensors, labels, or vertices is indicated below each significance map. Sensor-level significance was assessed using cluster-based permutation paired t-tests (5,000 permutations, *p* < .05, two-tailed). Parcellation-level significance was assessed using paired t-tests with FDR correction (*p* < .05). Vertex-level significance was assessed using spatial cluster-based permutation tests (1,024 permutations, *p* < .05, family-wise error corrected). Right hemisphere results are shown in Supplementary Figure S3.

At the source level, parcellation-based analysis with FDR correction identified 12 of 68 Desikan–Killiany labels (17.6%) with significant condition differences (minimum FDR-corrected *p* < .001). All significant labels showed higher exponents during rest than during Inscapes and were predominantly located in bilateral parietal regions including the superior parietal cortex, precuneus, inferior parietal cortex, and paracentral lobule, as well as the posterior cingulate cortex and precentral gyrus.

Vertex-level cluster-based permutation analyses corroborated and extended these findings, revealing two clusters encompassing 14,853 vertices (72.5% of the cortical surface; t-max = 7.14, *p* < .001), with higher exponent values during rest distributed across bilateral parietal, precuneus, and sensorimotor regions. Although the bilateral pericalcarine cortex showed a numerically higher exponents during Inscapes than during rest (pericalcarine-lh: rest = 1.01, Inscapes = 1.07; pericalcarine-rh: rest = 1.02, Inscapes = 1.08), visible as negative Cohen’s d values in the medial view of Figure 1, this effect did not survive FDR correction at the parcellation level (FDR-corrected *p* = .075 and .102, respectively). Representative power spectral densities with spectral parameterization model fits at frontal, central, parietal, and occipital sensors are provided in Supplementary Figures S1–S2.

### Aperiodic Offset

Aperiodic offsets were examined using the same multi-level approach (Figure 2; right hemisphere in Supplementary Figure S4). At the sensor level, magnetometers showed a positive cluster across 85 sensors (cluster statistic = 301.30, *p* < .001), reflecting higher aperiodic offset values during eyes open rest than during Inscapes. Gradiometers similarly revealed a positive cluster of 135 sensors (cluster statistic = 496.83, *p* < .001), with higher offset values during rest across a widespread scalp distribution. Cohen’s d topographies for both sensor types showed spatially diffuse moderate effect sizes, consistent with a global shift in broadband power between conditions.

**Figure 2.**
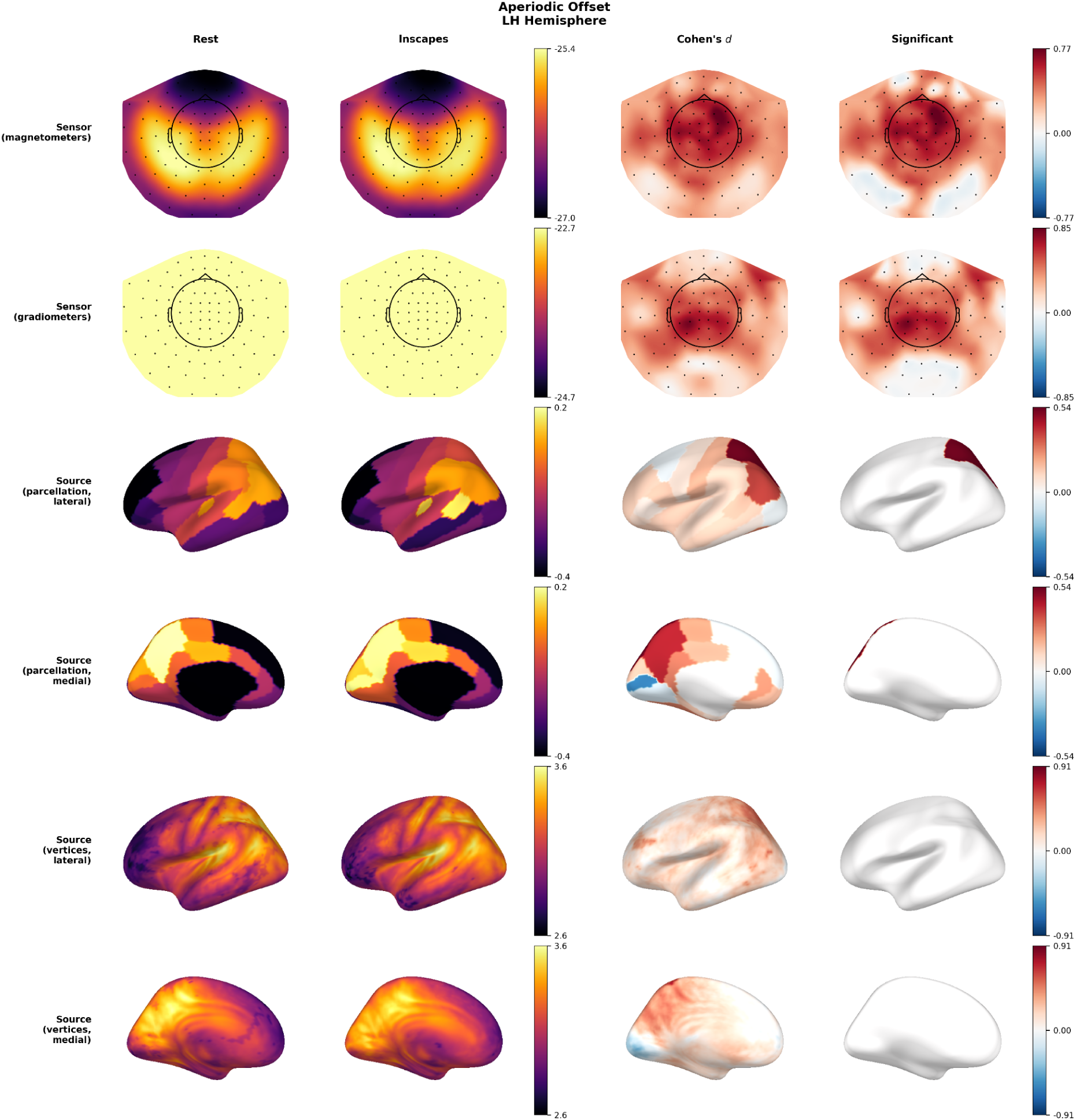
Multi level comparison of aperiodic offsets between eyes open rest and Inscapes, left hemisphere. Layout and conventions are the same as in Figure 1. Right hemisphere results are shown in Supplementary Figure S4.

At the source level, FDR-corrected parcellation analysis identified 2 of 68 labels (2.9%) with significant condition differences (minimum FDR-corrected *p* = .015), limited to the bilateral superior parietal cortex, both showing higher offset values during rest. Vertex-level analyses revealed one cluster totalling 2,196 vertices (10.7%; *t*-max = 6.68, *p* = .041), concentrated in parietal and posterior cingulate regions. Although the spatial extent of significant vertices was smaller for the offset than for the exponent, the topographic distribution of Cohen’s d values was broadly consistent between the two aperiodic measures.

### Alpha Power

Alpha band power was compared across all analysis levels (Figure 3; right hemisphere in Supplementary Figure S5). At the sensor level, magnetometers showed a positive cluster encompassing 51 sensors (cluster statistic = 300.36, *p* < .001), confirming greater alpha power during eyes open rest than during Inscapes, predominantly over occipital and parietal channels. Gradiometers revealed the most spatially extensive effect among all spectral parameters, with a positive cluster of 86 sensors (cluster statistic = 514.50, *p* = .001). Cohen’s d maps showed the largest effect sizes over posterior sensors, consistent with the well-established posterior distribution of alpha desynchronization during visual stimulation.

**Figure 3.**
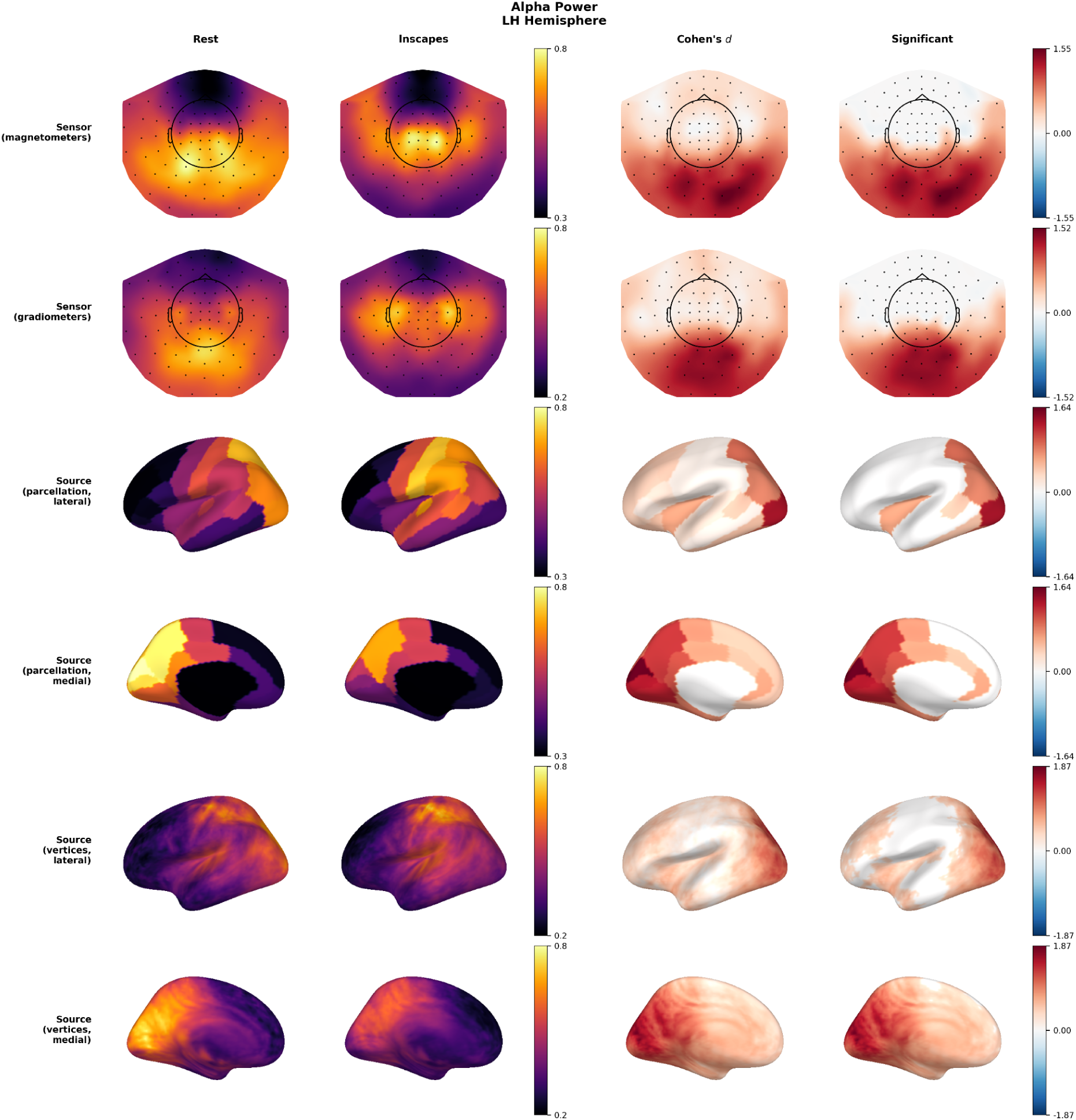
Multi level comparison of alpha power between eyes open rest and Inscapes, left hemisphere. Layout and conventions are the same as in Figure 1. Right hemisphere results are shown in Supplementary Figure S5.

At the source level, alpha power differences were the most widespread of all three spectral parameters. FDR-corrected parcellation analysis identified 38 of 68 labels (55.9%) with greater alpha power during rest than during Inscapes (minimum FDR-corrected *p* < .001), spanning bilateral occipital, parietal, temporal, and frontal regions. Vertex-level analyses yielded the most extensive spatial coverage of all measures, with two clusters comprising 15,309 vertices (74.7%; *t*-max = 10.75, *p* < .001), all reflecting greater alpha power during rest and spanning occipital, parietal, and temporal cortices. The Cohen’s d maps at the vertex level confirmed large effect sizes throughout the posterior cortex, tapering to moderate values anteriorly.

## Discussion

The present study used spectral parameterisation of MEG data to compare aperiodic neural activity between eyes open rest and Inscapes viewing in healthy adults. The central finding was that aperiodic exponents and offsets were higher during rest than during Inscapes across widespread frontoparietal and sensorimotor regions at both sensor and source levels. Alpha power was similarly greater during rest. These results demonstrate that Inscapes viewing produces measurable shifts in the temporal dynamics of neural activity that extend beyond the functional connectivity and averaged spectral power differences reported in prior work (Vanderwal et al., 2015, 2017; Vandewouw et al., 2021). The consistency of effects across sensor types, parcellation labels, and vertex level analyses strengthens the conclusion that Inscapes and eyes open rest engage distinct spectral regimes.

The steeper aperiodic exponent observed during rest relative to Inscapes can potentially be interpreted within the framework linking the spectral exponent to excitation/inhibition (E/I) balance (Gao et al., 2017). According to this framework, steeper power spectral slopes (higher exponents) reflect a shift toward greater inhibition, whereas flatter slopes (lower exponents) indicate a shift toward excitation. The present pattern, in which Inscapes produced flatter exponents across most of the cortex, is consistent with sustained visual stimulation increasing cortical excitability relative to the fixation cross condition. This interpretation aligns with evidence that aperiodic exponents decrease during wakefulness compared with sleep and anesthesia (Lendner et al., 2020) and that sensory engagement modulates the spectral slope in a modality specific manner (Waschke et al., 2021). However, it should be noted that the relationship between the aperiodic exponent and E/I balance remains a topic of active investigation. Recent pharmacological work has shown that the exponent does not universally track E/I ratio under all manipulations (Salvatore et al., 2024), and computational modeling suggests that multiple biophysical factors beyond synaptic E/I balance, including oscillatory relaxation processes (Muthukumaraswamy & Liley, 2018), can shape the aperiodic spectrum. The present findings are therefore best characterized as reflecting a shift in the broadband spectral regime that is consistent with, but not exclusively attributable to, changes in E/I balance.

The widespread reduction in alpha power during Inscapes viewing is consistent with the well established phenomenon of alpha desynchronisation during visual stimulation. Alpha oscillations are typically strongest during relaxed wakefulness without dynamic visual processing and are suppressed when external visual input engages cortical circuits. The present results extend this principle to a low demand, abstract visual paradigm and suggest that even minimal dynamic visual engagement is sufficient to attenuate alpha power relative to a static visual input (e.g., a fixation cross). Importantly, the spectral parameterisation approach allowed these periodic changes to be dissociated from the simultaneously occurring aperiodic changes, demonstrating that Inscapes modulates both oscillatory and broadband components of the power spectrum.

Although the pericalcarine cortex, corresponding to primary visual cortex (V1), did not show statistically significant condition differences when correcting for multiple comparisons, a consistent trend toward higher aperiodic exponents and offsets during Inscapes than during rest was observed across both parcellation and vertex levels. This pattern was opposite to the direction seen across the rest of the cortex and was visible as negative Cohen’s d values in the medial views of Figures 1 and 2. This directional trend is interpretively coherent: primary visual cortex is the region most directly engaged by the continuous visual input provided by Inscapes. If steeper aperiodic slopes index greater inhibitory tone, the tendency toward increased exponents in V1 during Inscapes may reflect enhanced feedforward inhibitory processing recruited during sustained visual stimulation, consistent with evidence that greater visual input intensity is followed by proportionally greater functional inhibition in visual areas (Manyukhina et al., 2024). While this observation should be interpreted cautiously given the lack of statistical significance, it suggests that the neural effects of Inscapes may not be uniform across the cortex and that the direction of aperiodic modulation could depend on whether a region is directly engaged by the stimulus. Such possibilities could merit direct follow-up in future studies.

To verify that the present findings are not contingent on the choice of spectral parameterisation algorithm, we conducted a supplementary analysis using Irregular-Resampling Auto-Spectral Analysis (IRASA; Wen & Liu, 2016), which separates fractal and oscillatory spectral components through a resampling-based approach that does not rely on explicit parametric modeling of spectral peaks (see Supplementary Figures S6–S7). IRASA replicated all primary sensor level findings: aperiodic exponents and offsets were higher during rest than during Inscapes across nearly all gradiometers (198/204 and 193/204 sensors, respectively) and magnetometers (102/102 sensors for both measures), with cluster-corrected *p* values < .001 for all four tests. Alpha power was greater during rest in posterior sensors, consistent with the spectral parameterisation results. The convergence of IRASA and spectral parameterisation on all primary findings, despite their different assumptions for separating periodic and aperiodic components (Gerster et al., 2022), strengthens confidence in the robustness of the present results.

The present findings have direct practical implications for the growing use of Inscapes as a substitute for eyes open rest in neuroimaging research. Although Inscapes has been shown to preserve functional connectivity patterns similar to rest (Vanderwal et al., 2015, 2017) and to reduce head motion (Vandewouw et al., 2021), the current results demonstrate that the two conditions are not equivalent in terms of aperiodic spectral dynamics. Studies that use Inscapes to acquire data for analyses involving broadband spectral power, spectral slope, or E/I balance estimates should recognise that Inscapes introduces systematic shifts in these parameters relative to rest. This does not diminish the value of Inscapes as a tool for improving data quality, particularly in pediatric and clinical populations where compliance is a primary concern, but it does caution against treating Inscapes and rest as directly interchangeable for all analytical purposes. Future work could consider reporting both conditions where feasible, or explicitly accounting for the known spectral differences when comparing across studies that use different baseline paradigms.

Several limitations should be acknowledged. Firstly, the sample comprised healthy young adults so the generalisability of these findings to other populations remains to be established. Developmental changes in the aperiodic exponent may interact with condition effects in ways not captured here and different outcomes may be seen in, for example, children or elderly participants (Hill et al., 2022; Voytek et al., 2015).

Secondly, the study did not include behavioural measures of drowsiness or eye tracking data, so it is not possible to determine whether differences in arousal or eye movement patterns contributed to the observed spectral differences. Thirdly, the condition order was not counterbalanced across participants, which may introduce order effects. Fourthly, the effect of Inscapes viewing on head motion was not measured and so potential interactions of any such difference and parameter changes between conditions may be present. Prior work has shown that the main influence of head motion on the MEG signal occurs in the range below 0.5 Hz and that filtering this range out of the signal minimises the influence of head motion on MEG time series (Messaritaki et al., 2017). We therefore repeated our analysis while band-pass filtering between 0.5-55 Hz. Results remain the same, suggesting that head motion itself is not driving the rest-Inscapes differences (Supplementary Figures S8-13). Finally, the present study presented only the visual component of Inscapes without the accompanying audio track, which differs from the original paradigm design (Vanderwal et al., 2015). It is possible that the inclusion of the pentatonic piano score would further modulate aperiodic activity, and the present results should therefore be interpreted as reflecting the effects of the visual component alone.

In conclusion, this study provides evidence that Inscapes viewing and eyes open rest produce distinct aperiodic spectral profiles, as measured by MEG. The condition differences in aperiodic exponent, offset, and alpha power were robust across sensor types, cortical parcellations, and vertex level analyses. These findings indicate that the visual aspect of Inscapes, despite preserving many features of resting state connectivity, engages a different broadband spectral regime that may reflect increased cortical excitability during visual stimulation. These results underscore the importance of spectral parameterisation as a complementary tool for characterising neural state differences and suggest that researchers employing Inscapes should consider how its effects on aperiodic dynamics may influence downstream analyses.

## Acknowledgements

The authors thank all participants for their time and effort. This work was supported by funding from the National Science and Technology Council (111-2410-H-038-009-MY2, NSTC 113-2410-H-008-082, NSTC 114-2410-H-008-066) and the Taiwan Ministry of Education Higher Education Sprout Project to TYH and the National Science and Technology Council (NSTC113-2423-H-038-002-MY3; NSTC114-2410-H-038-037) to NWD. In addition, we would like to thank the Imaging Center for Integrated Body, Mind and Culture Research at NTU for MRI and MEG data collection.

## Author contributions

Tzu-Yu Hsu: Conceptualisation, Funding, Investigation, Methodology, Data Curation, Software, Visualisation, Writing- Original draft preparation, Supervision Ko-Ping Chou: Investigation Yi-Ju Liu: Investigation Niall W. Duncan: Conceptualisation, Methodology, Writing- Original draft preparation, Supervision

## Data and code availability

The data and code required to reproduce the results presented in this work are available at

## Supplementary Material

### S1. Power Spectral Density at Representative Sensors

To provide visual confirmation that the spectral parameterization algorithm adequately captured the shape of the power spectra, Figures S1 and S2 display power spectral densities at four representative sensors spanning frontal, central, parietal, and occipital regions. Representative sensors were selected by dividing the anterior–posterior axis of sensor positions into four percentile-based zones (occipital: 0–15th percentile; parietal: 15–40th; central: 40–60th; frontal: 75–100th) and choosing the sensor closest to the midline within each zone. For each sensor, individual-subject PSDs (light traces) and the group-mean PSD (bold trace) are shown alongside the spectral parameterization full model fit (dashed line) and the aperiodic component fit (dotted line). The legend indicates the number of subjects contributing to each condition and the group-mean aperiodic exponent (β ± SD).

Power spectra were estimated using Welch’s method with 50% overlap on 4-second epochs, over a frequency range of 5–55 Hz. These plots illustrate that the 1/f model provided a close fit to the empirical spectra across all regions and that the alpha peak was clearly resolved above the aperiodic background in both conditions.

**Figure S1.**
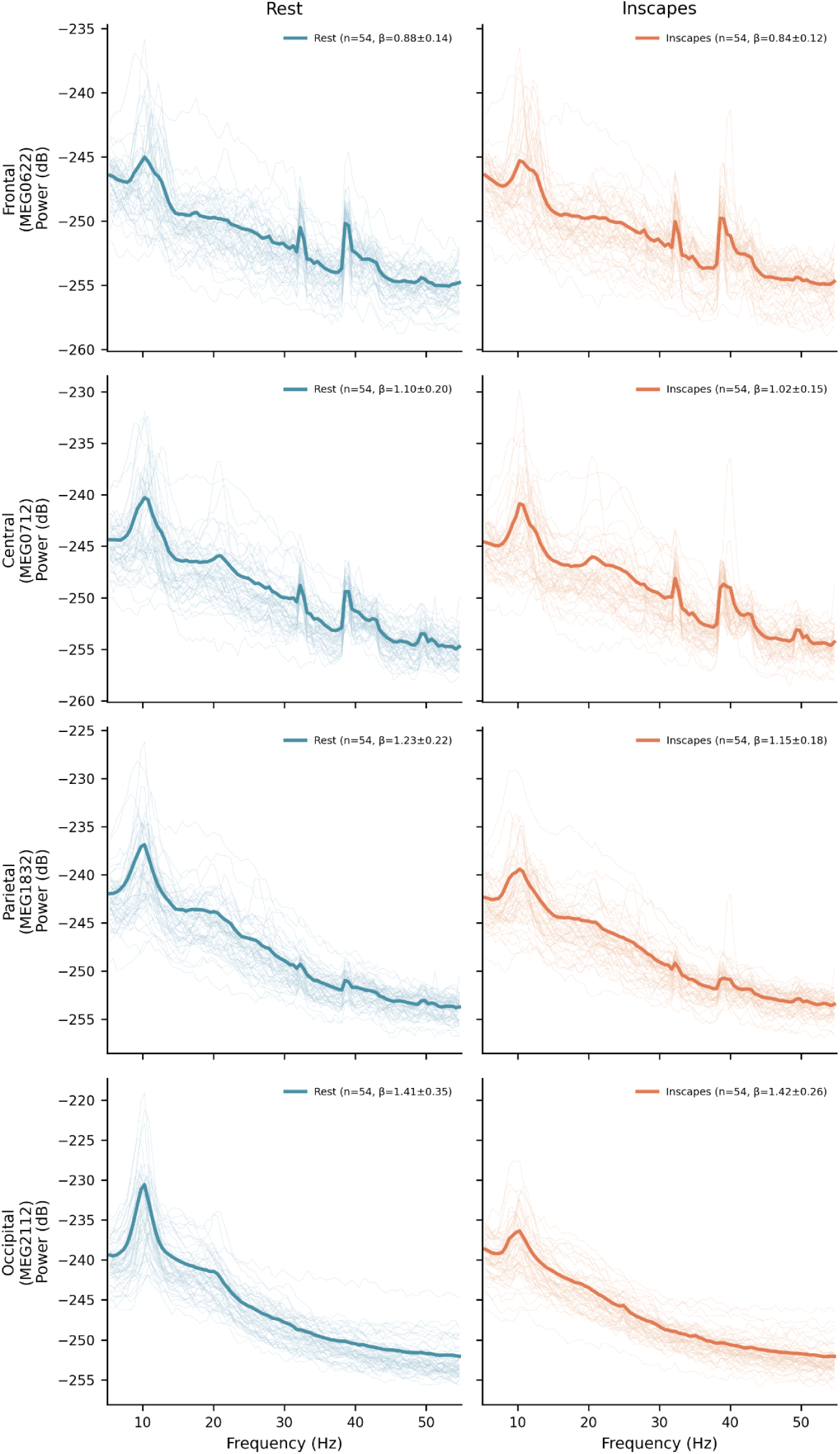
Power spectral densities at representative gradiometer sensors. Each row corresponds to a cortical region (frontal, central, parietal, occipital), with separate columns for the rest (left) and Inscapes (right) conditions. Light traces represent individual subject spectra; bold traces represent the group mean. The dashed line shows the spectral parameterization full model fit and the dotted line shows the aperiodic component fit applied to the group mean spectrum. The aperiodic exponent (β ± SD) is displayed in each legend.

**Figure S2.**
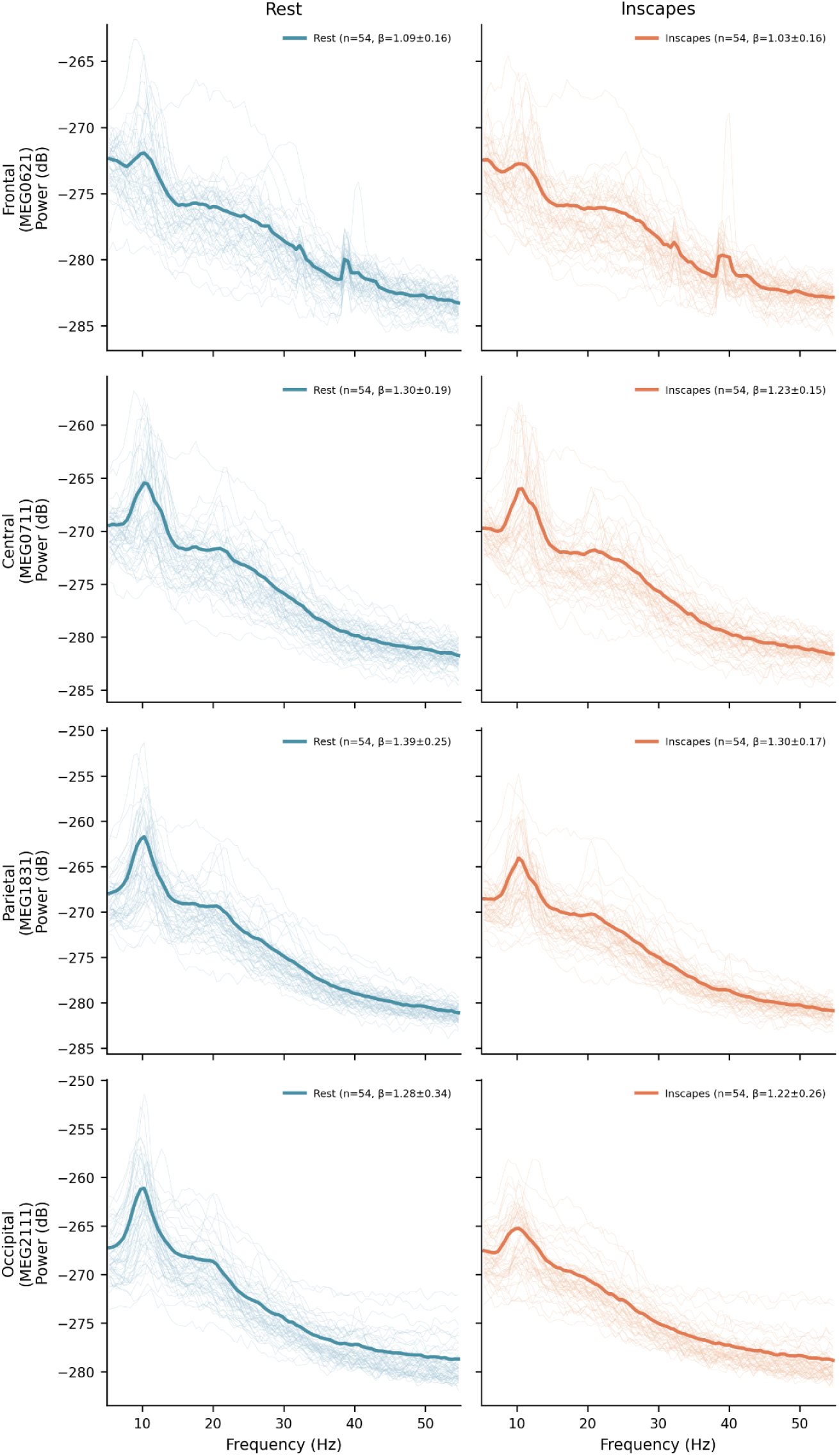
Power spectral densities at representative magnetometer sensors. Layout and conventions are the same as in Figure S1.

### S2. Right Hemisphere Source-Level Results

The main manuscript presents sensor-level and source-level results for the left hemisphere (Figures 1-3). To demonstrate that the condition effects were bilateral, Figures S3–S5 display the corresponding right hemisphere results for exponent, offset, and alpha power. The spatial distribution of condition differences in the right hemisphere closely mirrored that of the left hemisphere across all three spectral parameters, confirming that the effects reported in the main text were not lateralized.

**Figure S3.**
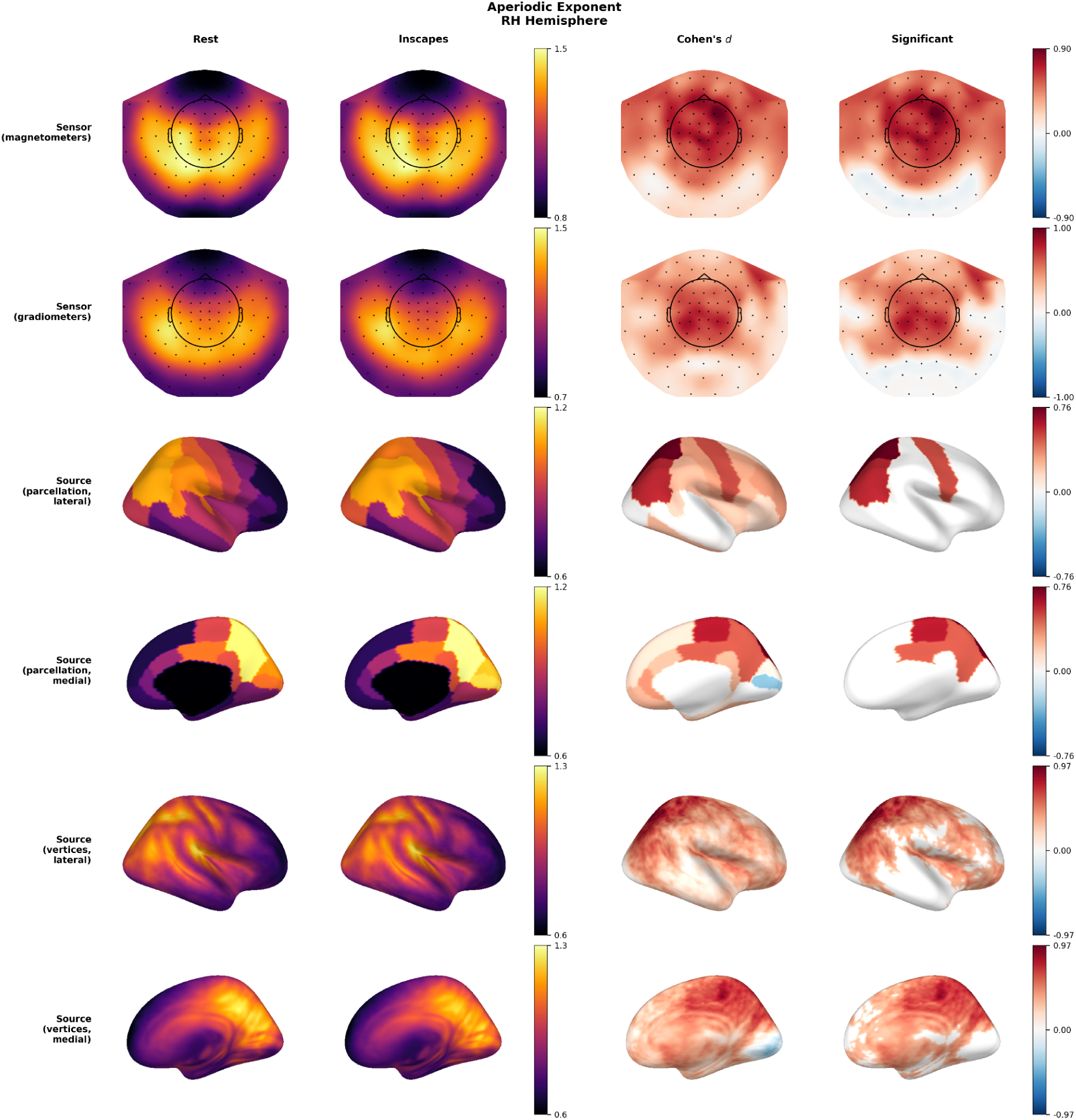
Multi level comparison of aperiodic exponents, right hemisphere. Layout and conventions are the same as in Figure 1.

**Figure S4.**
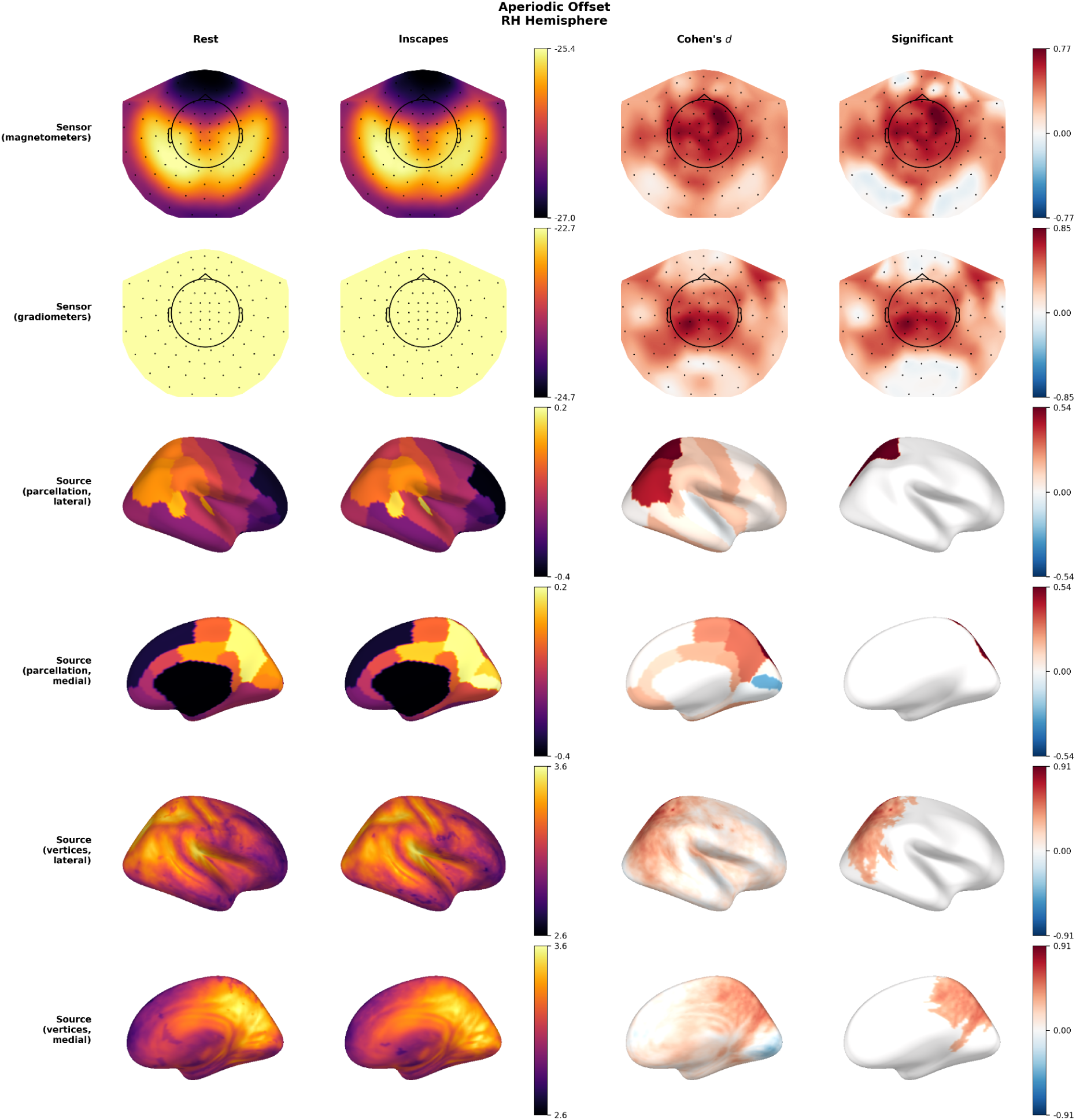
Multi level comparison of aperiodic offsets, right hemisphere. Layout and conventions are the same as in Figure 1.

**Figure S5.**
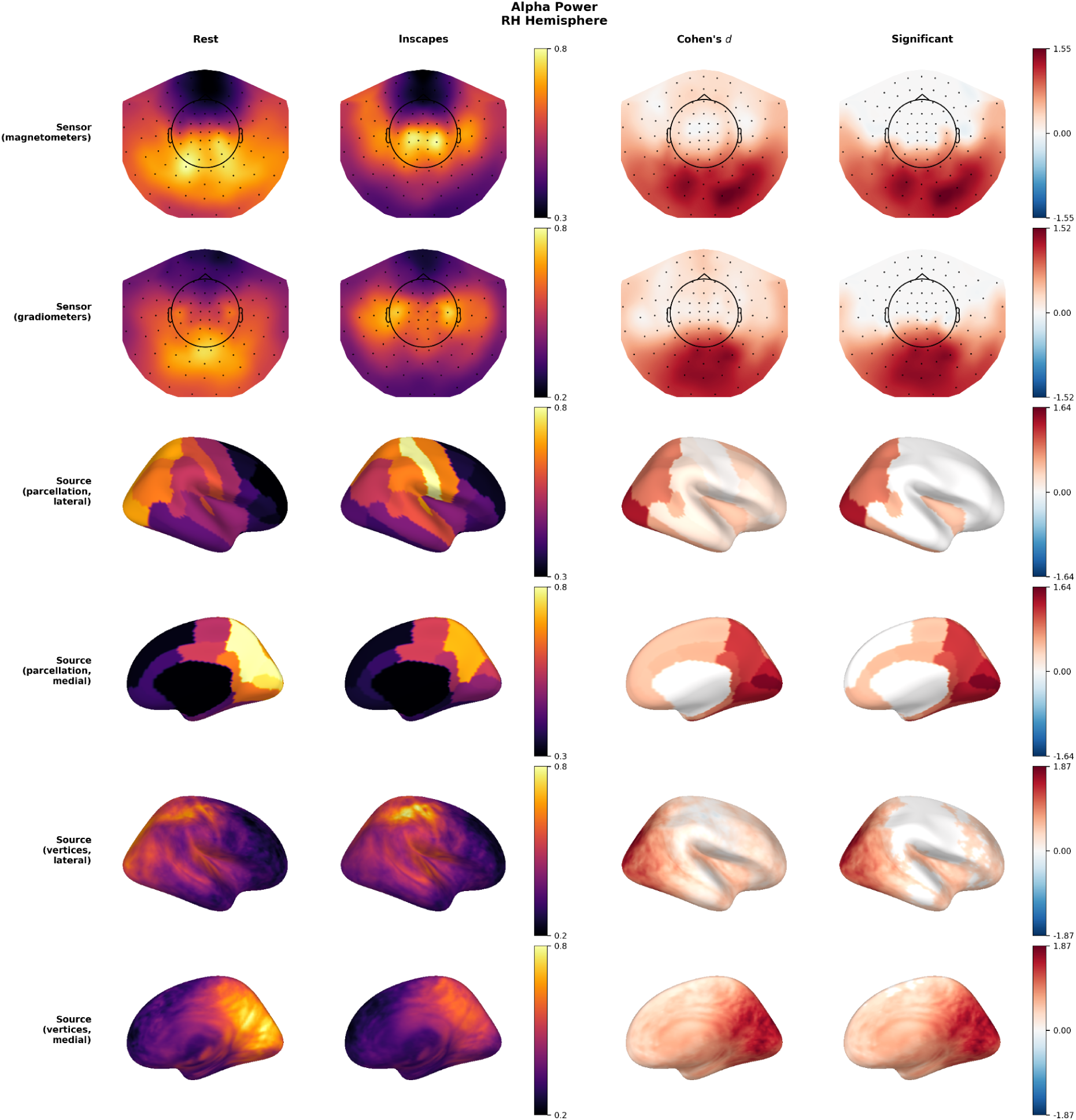
Multi level comparison of alpha power, right hemisphere. Layout and conventions are the same as in Figure 1.

### S3. Supplementary Methods: IRASA Analysis

To assess whether the condition differences in aperiodic parameters were dependent on the choice of spectral decomposition method, we conducted a supplementary sensor-level analysis using Irregular-Resampling Auto-Spectral Analysis (IRASA; Wen & Liu, 2016) as implemented in the YASA package (Vallat & Walker, 2021).

Unlike spectral parameterization (specparam), which models periodic peaks as Gaussians superimposed on a parametric aperiodic fit, IRASA separates fractal and oscillatory components by exploiting the scale-free property of aperiodic signals.

Specifically, the algorithm resamples the time series by a set of non-integer factors (h = 1.1 to 1.9 in increments of 0.05, yielding 17 resampling factor pairs), computes the geometric mean of the auto-power spectra for each up/down-sampled pair, and takes the median across all pairs to extract the fractal (aperiodic) component. The oscillatory component is then obtained by subtracting the fractal estimate from the original power spectrum.

IRASA was applied to the same preprocessed continuous MEG data used in the main analysis. For each participant and condition, data were segmented from artifact-free annotations, and IRASA was computed per sensor using a 4-second sliding window within a frequency band of 5–50 Hz. The aperiodic exponent and offset were estimated by fitting a linear regression to the fractal power spectrum in log-log space across the 5–50 Hz range. Alpha peak frequency and power were extracted from the oscillatory residual spectrum within the 8–13 Hz band by identifying the frequency with the maximum oscillatory power. Sensors with fewer than 10 valid frequency bins or with a positive slope (indicating a poor fractal fit) were excluded from analysis.

Statistical analysis mirrored the main sensor-level approach. For each metric, a paired t-test was computed at every sensor, and cluster-based permutation testing (5,000 permutations, cluster-forming threshold p < .05) was used to identify spatially contiguous sensor clusters with significant condition differences while controlling the family-wise error rate.

### S4. Supplementary Results: IRASA

IRASA replicated the primary aperiodic findings from the main spectral parameterization analysis. For gradiometers (Figure S6), aperiodic exponents were significantly higher during rest than during Inscapes in a single large cluster encompassing 198 of 204 sensors (cluster statistic = 1018.01, p < .001), with mean exponent values of 1.35 during rest and 1.25 during Inscapes. Aperiodic offsets showed an analogous pattern across 193 of 204 sensors (cluster statistic = 984.68, p < .001). Alpha power and alpha peak frequency did not show significant condition differences at the cluster-corrected level.

**Figure S6.**
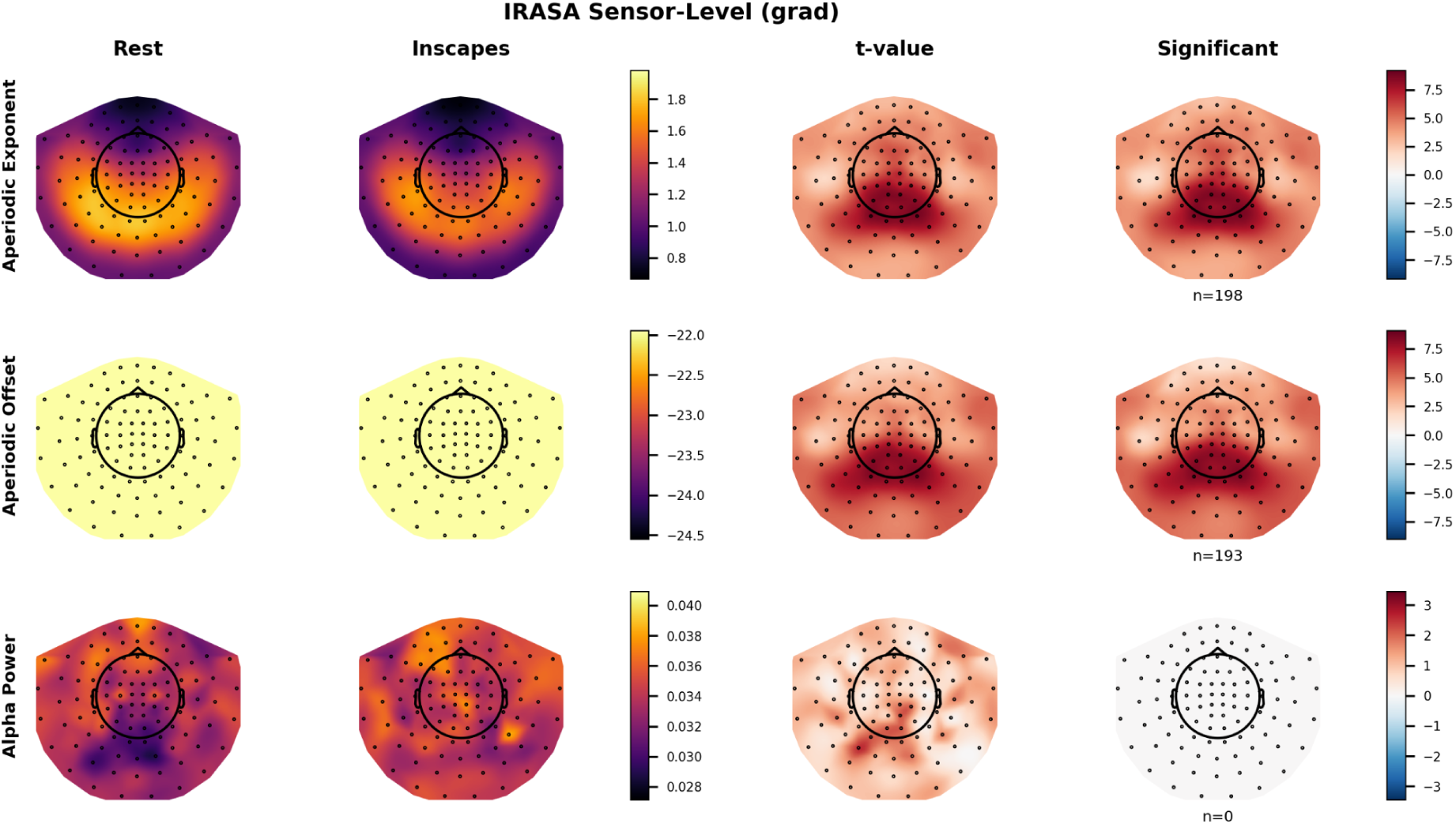
IRASA sensor-level results for gradiometers. Topographical maps of aperiodic exponent and offset for the rest condition (left), Inscapes condition (middle), and their statistical contrast (right). Sensors belonging to significant clusters (p < .05, cluster-corrected) are highlighted.

For magnetometers (Figure S7), IRASA yielded consistent results for aperiodic measures. Aperiodic exponents were higher during rest across all 102 sensors (cluster statistic = 614.23, p < .001; rest mean = 1.47, Inscapes mean = 1.34), and offsets were similarly higher during rest across all 102 sensors (cluster statistic = 580.86, p < .001). As with gradiometers, neither alpha power nor alpha peak frequency showed significant condition differences.

**Figure S7.**
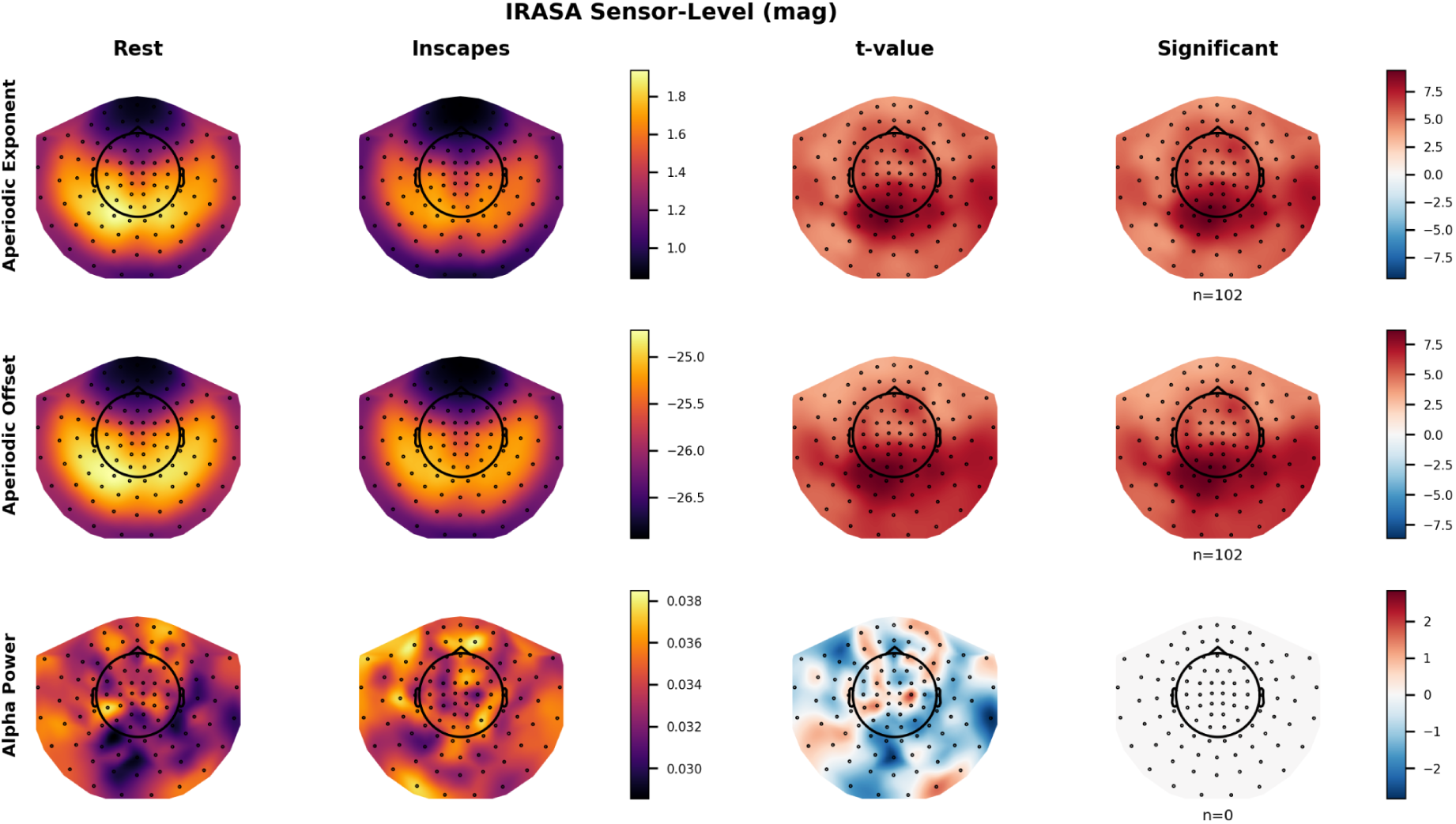
IRASA sensor-level results for magnetometers. Topographical maps of aperiodic exponent and offset for the rest condition (left), Inscapes condition (middle), and their statistical contrast (right). Sensors belonging to significant clusters (p < .05, cluster-corrected) are highlighted.

### S5. Comparison Between IRASA and Spectral Parameterization

The IRASA and spectral parameterization analyses converged on the primary aperiodic findings: both methods identified significantly higher aperiodic exponents and offsets during rest compared with Inscapes. However, the two methods diverged on periodic measures: spectral parameterization detected significantly greater alpha power during rest, whereas IRASA did not find significant alpha power differences at the cluster-corrected level. The two methods differ in their approach to separating periodic and aperiodic spectral components. Spectral parameterization fits Gaussian peaks to the power spectrum and estimates the aperiodic component from the residual, whereas IRASA uses a resampling-based strategy that leverages the scale-invariance of fractal processes to extract the aperiodic component without explicitly modeling oscillatory peaks. This difference means that IRASA’s resampling procedure may absorb some of the alpha-band power variation into the fractal component, effectively redistributing what spectral parameterization assigns to the periodic peak. The convergence on aperiodic findings provides strong evidence that the condition differences in aperiodic parameters reflect genuine neural phenomena rather than method-specific artifacts.

## S6. Robustness Check: High-Pass Filter at 0.5 Hz

To assess whether the main findings were sensitive to the choice of high-pass filter, we repeated the full spectral parameterization analysis with data filtered between 0.5 and 55 Hz, rather than the 0.1–55 Hz bandpass used in the main analysis. A higher high-pass cutoff more aggressively attenuates slow drifts and head-movement artifacts that could influence the aperiodic fit, providing a test of whether the main findings are robust to this preprocessing choice. Figures S8–S13 present the multi-level results for the aperiodic exponent, aperiodic offset, and alpha power, respectively, using the same layout as main Figures 1–3.

The results were nearly identical to the main analysis. At the sensor level, all significant clusters were replicated with comparable spatial extent and effect sizes: magnetometer exponents spanned 88 sensors (cluster statistic = 367.49, p < .001), offsets 86 sensors (303.44, p < .001), and alpha power 50 sensors (301.39, p < .001); gradiometer exponents spanned 138 sensors (549.68, p < .001), offsets 132 sensors (489.44, p < .001), and alpha power 86 sensors (514.10, p = .001). At the parcellation level, FDR-corrected analysis identified 14 of 68 labels for exponents (compared with 12 in the main analysis), 2 of 68 for offsets (identical), and 38 of 68 for alpha power (identical). The close correspondence between the two filter settings indicates that the reported condition differences are not driven by low-frequency artifacts associated with the high-pass filter cutoff and are robust to this analytical choice.

**Figure S8.**
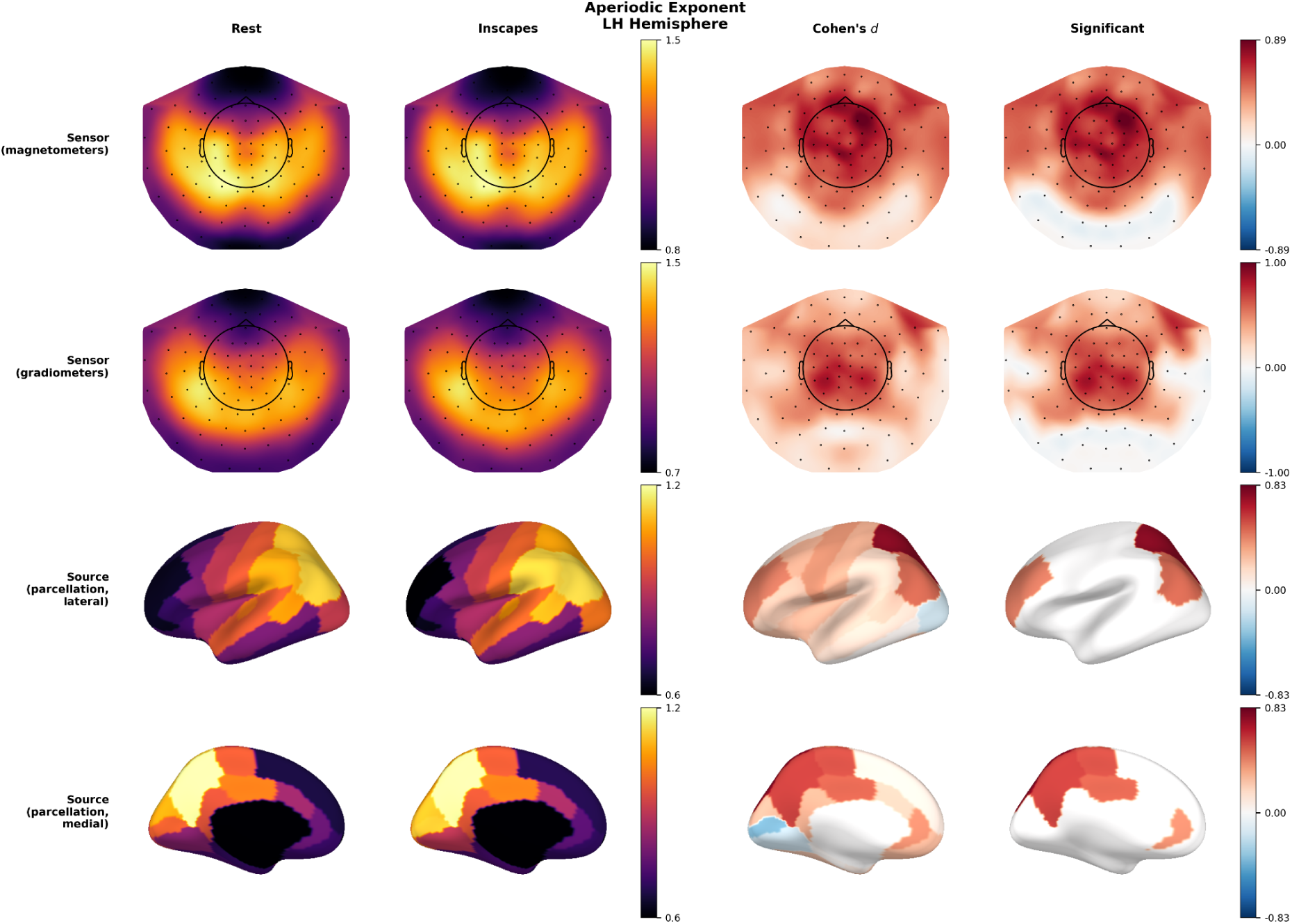
Multi level comparison of aperiodic exponent between eyes open rest and Inscapes using 0.5-55 Hz filtering data (left hemisphere). Layout and conventions are the same as in main Figure 1.

**Figure S9.**
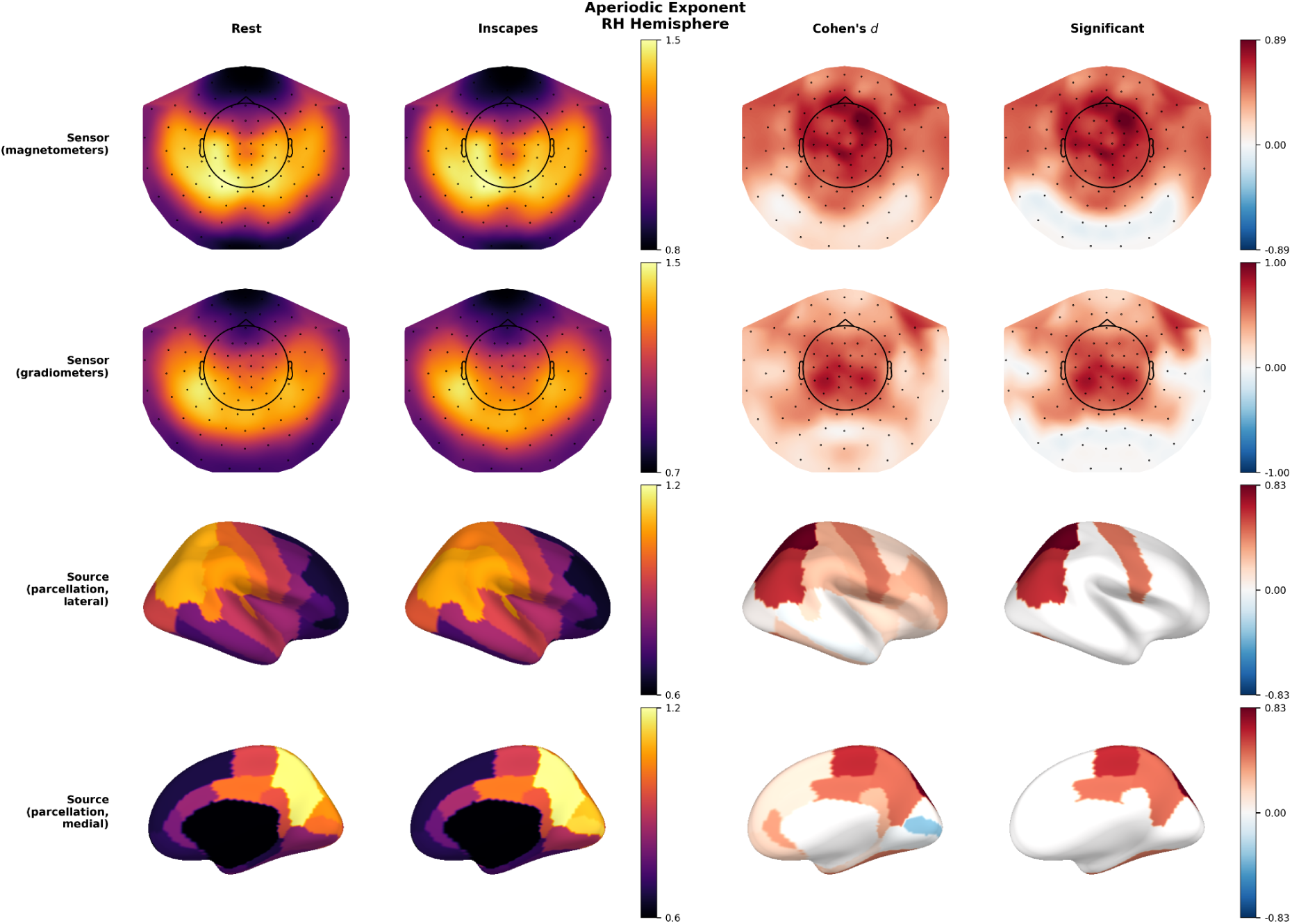
Multi level comparison of aperiodic exponent between eyes open rest and Inscapes using 0.5-55 Hz filtering data (right hemisphere). Layout and conventions are the same as in main Figure 1.

**Figure S10.**
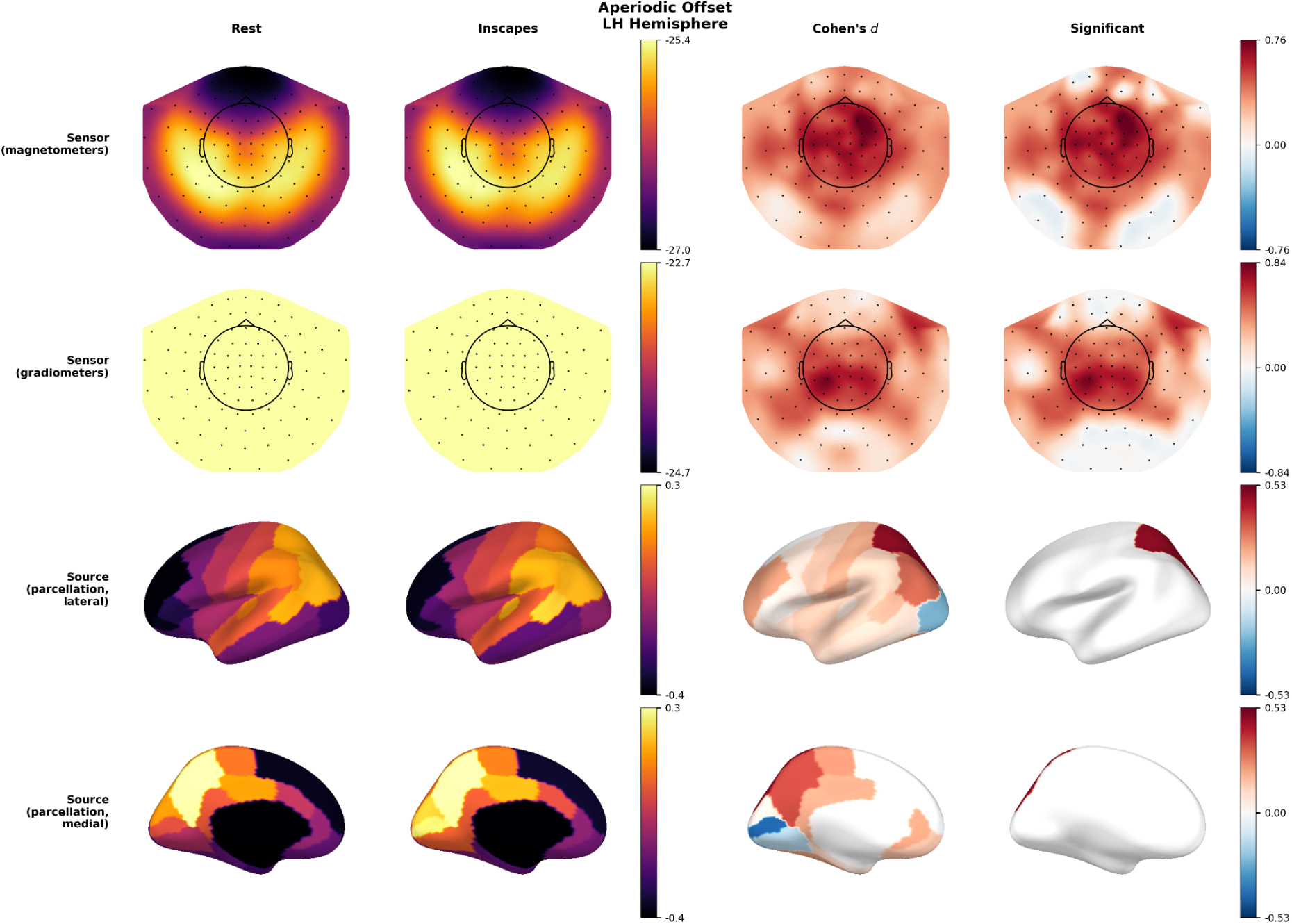
Multi-level comparison of aperiodic offset between eyes open rest and Inscapes using 0.5–55 Hz filtered data (left hemisphere). Layout and conventions are the same as in main Figure 1.

**Figure S11.**
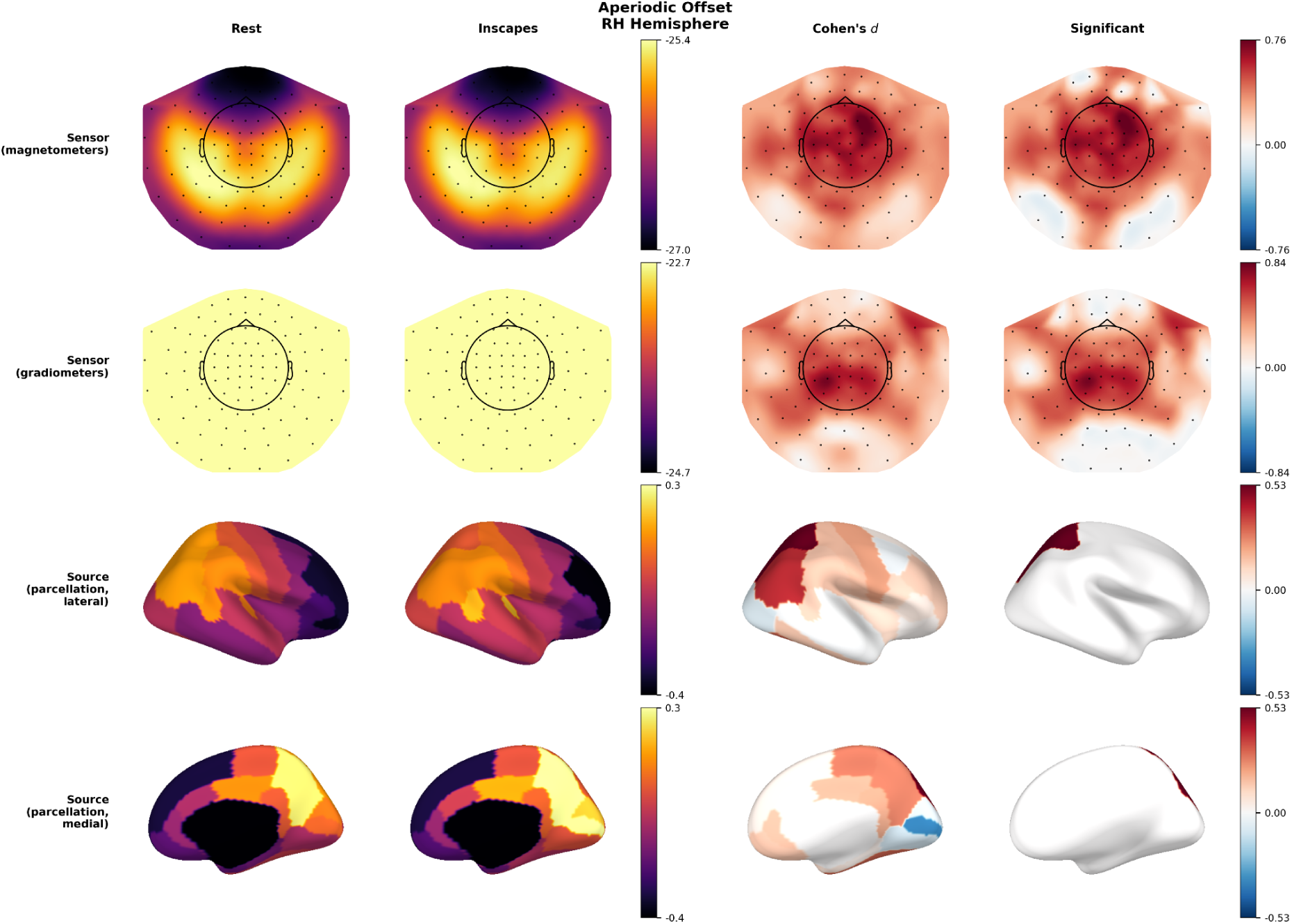
Multi-level comparison of aperiodic offset between eyes open rest and Inscapes using 0.5–55 Hz filtered data (right hemisphere). Layout and conventions are the same as in main Figure 1.

**Figure S12.**
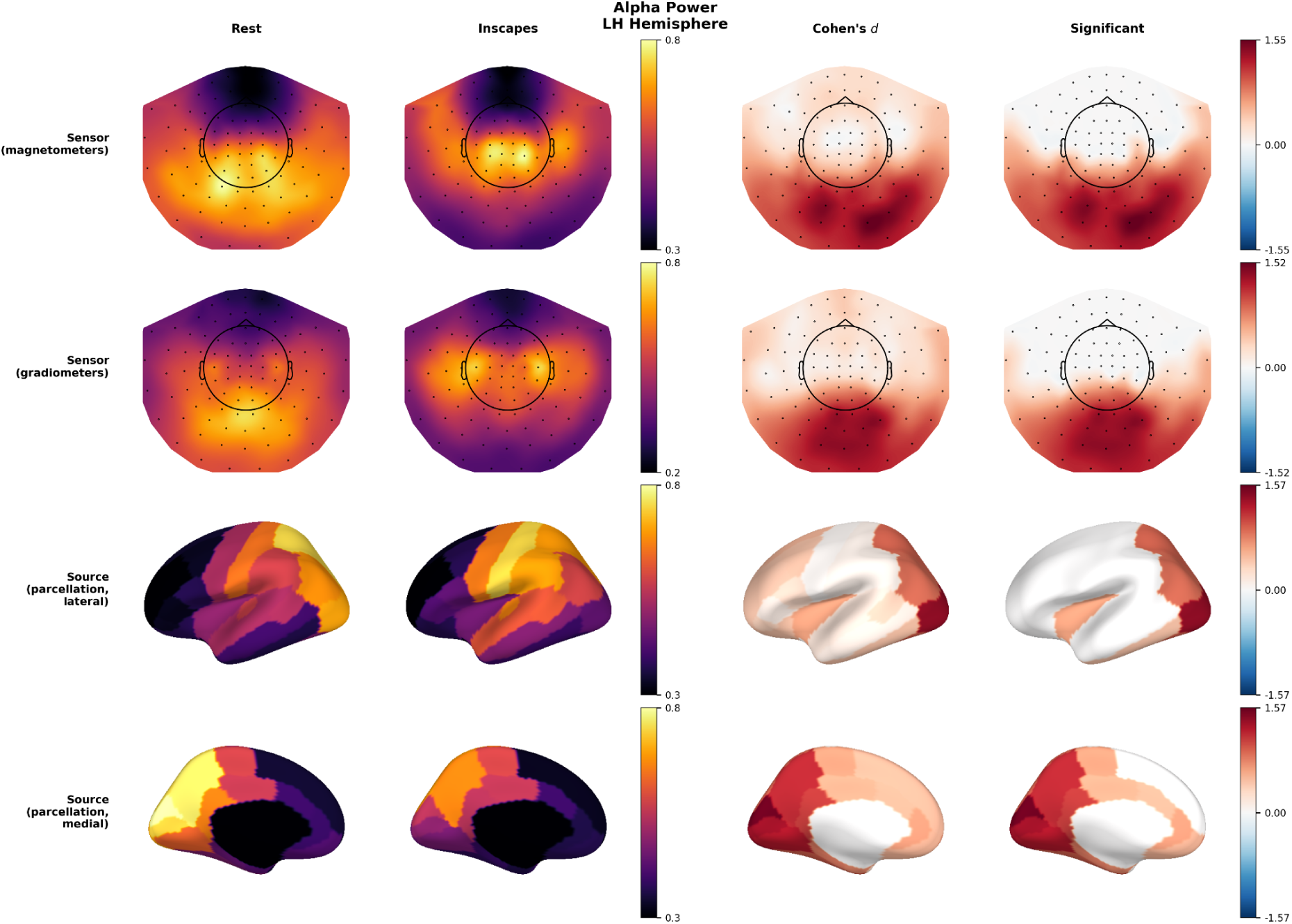
Multi-level comparison of alpha power between eyes open rest and Inscapes using 0.5–55 Hz filtered data (left hemisphere). Layout and conventions are the same as in main Figure 1.

**Figure S13.**
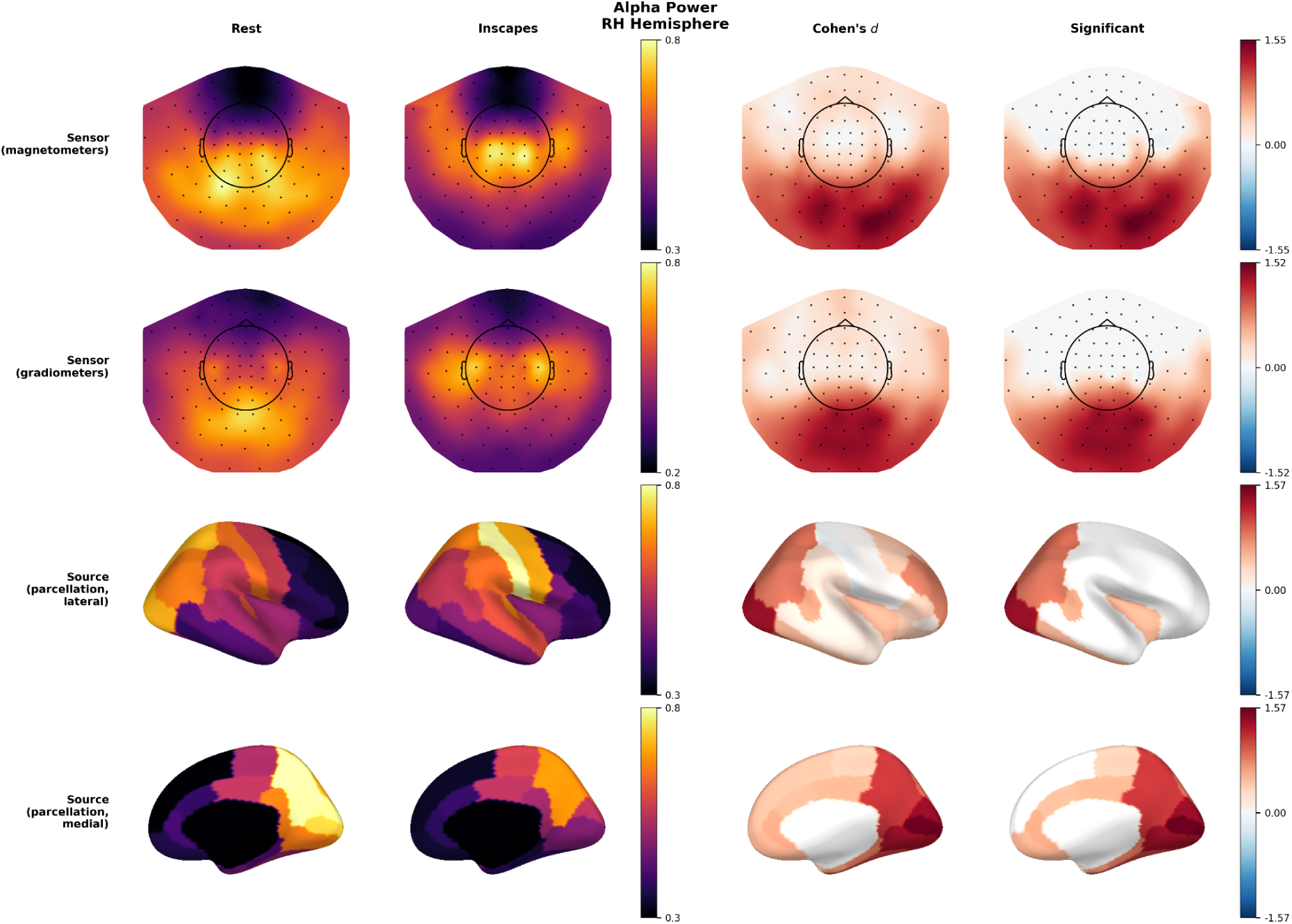
Multi-level comparison of alpha power between eyes open rest and Inscapes using 0.5–55 Hz filtered data (right hemisphere). Layout and conventions are the same as in main Figure 1.

